# Neuronal mechanism of the encoding of socially familiar faces in the striatum tail

**DOI:** 10.1101/2023.05.10.540108

**Authors:** Jun Kunimatsu, Hidetoshi Amita, Okihide Hikosaka

**Affiliations:** Laboratory of Sensorimotor Research, National Eye Institute, National Institutes of Health, Bethesda, MD 20892, USA; Division of Biomedical Science, Institute of Medicine, University of Tsukuba, Tsukuba, Ibaraki 305-8577, Japan; Transborder Medical Research Center, University of Tsukuba, Tsukuba, Ibaraki 305-8577, Japan; Systems Neuroscience Section, Center for the Evolutionary Origins of Human Behavior, Kyoto University, Inuyama, Aichi 484-8506, Japan

**Author notes:** Correspondence to:* Jun Kunimatsu, Ph.D. Division of Biomedical Science, Institute of Medicine, University of Tsukuba, Tsukuba, Ibaraki 305-8577, Japan.

## Abstract

Although we can quickly locate a familiar person even in a crowd, the underlying neuronal mechanism remains unclear. Recently, we found that the striatum tail (STRt), which is part of the basal ganglia, is sensitive to long-term reward history. Here, we show that long-term value-coding neurons are involved in the detection of socially familiar faces. Many STRt neurons respond to facial images, especially to those of socially familiar persons. Additionally, we found that these face-responsive neurons also encode the stable values of many objects based on long-term reward experiences. Interestingly, the strength of neuronal modulation of social familiarity bias (familiar or unfamiliar) and object value bias (high-valued or low-valued) were positively correlated. These results suggest that both social familiarity and stable object-value information are mediated by a common neuronal mechanism. This mechanism may contribute to the rapid detection of familiar faces in real-world contexts.

**Teaser:** The common mechanism mediating social familiarity and stable object-value information may contribute to rapid detection of familiar faces.

## Introduction

We can quickly find familiar people in a crowd. This search skill is critical, particularly for social animals, including humans. However, the underlying neuronal mechanism remains unclear. Facial information is one of the most important cues for identifying a person (*1*). Studies have suggested that there is a hierarchical organization of face processing along the posteroanterior axis of the temporal cortex in macaques (*2, 3*). The posterior area represents particular facial features (i.e., eye, nose, and lips), the middle area represents whole faces, and the anterior area represents identical faces, regardless of face orientation. In this manner, the face processing systems for identifying individuals have recently become clearer. However, most studies on face recognition systems have used unfamiliar faces. Therefore, it remains unclear how facial information is combined with social information (e.g., social familiarity) to identify individuals based on personal relationships.

Recently, our group reported that neurons in the striatum tail (STRt), including the caudate tail (CDt) and putamen tail, show different responses to fractal objects based on their associations with different reward sizes (*4, 5*). Interestingly, visual responses rarely changed across blocks in a reversal learning task; instead, visual responses gradually changed across days when each object was continuously associated with a fixed reward size (*4-8*). Macaque monkeys can learn the value of more than 500 objects and retain their value memories for more than 100 days (*6*). Monkeys can quickly distinguish high-valued objects among many objects and tend to look at high-valued objects or automatically avoid low-valued objects (*9, 10*). This automatic behavior is impaired by CDt inactivation (*7*). These results indicate that the STRt is involved in the rapid detection of valued objects based on long-term memory. Because the STRt receives dense inputs from the temporal cortex (TE and TEO), including from facial areas (*11-14*), it may also be involved in face detection. Previous studies have reported that patients in the early stage of Huntington’s disease, whose brains have distinct neuronal loss and gliosis in the CDt (*15*), showed deficits in face recognition and discrimination (*16*).

These results raise several questions. Does the STRt encode facial information? If so, does this represent a face’s social familiarity? Do the same neurons encode social familiarity and object values? We addressed these questions in the present study.

## Results

This study aimed to examine whether the STRt shows face selectivity and encodes both facial social familiarity and object value. For this purpose, we tested neuronal responses to facial images under two conditions: social familiarity and object value. In the social-familiarity condition, we used 12 images that included three categories: a) facial images of socially familiar persons who took daily care of the subject monkeys for more than 1 year [social familiarity (+)], b) facial images of socially unfamiliar persons [social familiarity (-)], and c) fractal objects associated wi
th no reward as control (Fig. 1A). Eight fractal objects were used in the object-value condition (Fig. 1B). The monkeys learned the values of these objects using a previously employed stable object-value task (Fig. S1A, (*4-7, 9, 17*). In each trial of the stable object-value task (Fig. S1C), one fractal object was randomly selected from a set of eight objects and presented as the target. The monkey made a saccade to the target followed by reward delivery. Each object was associated with a fixed amount of reward (small or large) across trials and sessions. Four of the eight objects were always associated with a large reward [high-valued objects, object value (+)], and the other four objects were always associated with a small reward [low-valued objects, object value (-)]. We then examined the monkey’s eye movements with a free-viewing task (Fig. 1C, left) from a few days to a few months after the stable object-value learning took place (Fig. S1B). In this task, four objects were randomly selected from a set of eight and presented simultaneously. The monkeys were then allowed to look at them for 2 s. A fixed amount of reward was delivered independent of the looking behavior or presented objects. Therefore, looking bias would reflect the long-term object-value memories. Figure 1C right panel illustrates the first saccade bias in the free-viewing task after several learning sessions (> five sessions). The monkeys mostly made first saccades to value (+) objects and avoided value (-) objects (p = 4.82 × 10-10, unpaired t-test). These results suggest that the monkeys had learned and retained stable object values and thereby found value (+) objects automatically.

**Fig. 1.**
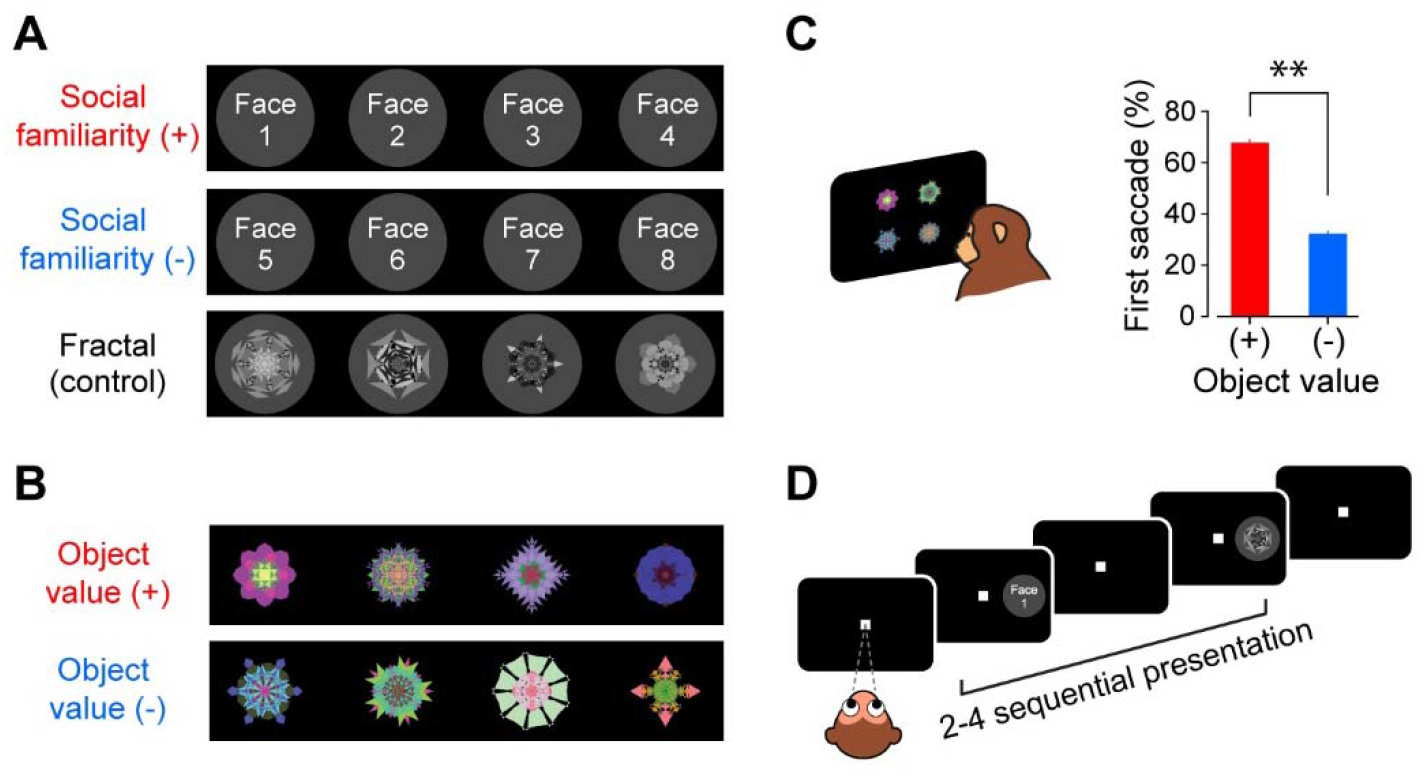
Behavioral and neuronal tests for social familiarity and stable object value. **(A)** In the social-familiarity condition, facial images were divided into the following three categories: (1) socially familiar persons’ faces [social familiarity (+), top], (2) socially unfamiliar persons’ faces [socially familiarity (-), middle], and (3) fractal objects associated with no reward (bottom). Face photographs were omitted. **(B)** In the stable object-value condition, facial images were divided into the following two groups: (1) fractal objects consistently associated with a large reward [object value (+), top] and (2) fractal objects consistently associated with a small reward [object value (-), bottom]. **(C)** During the free-viewing task, four fractal objects were simultaneously presented (left). The monkeys were free to look at these objects for 2 s without feedback. The first saccade to value (+) or value (-) objects in 94 sessions of the free-viewing task (right; mean ± SEM; three monkeys). **(D)** In both the social-familiarity and stable object-value conditions, facial images were sequentially presented in the neuron’s receptive field as the monkey was fixating on the center (passive-viewing task).

We tested whether STRt neurons responded to facial and fractal images using a passive-viewing task (Fig. 1A). In this task, each object was presented at the neuron’s receptive field while the monkey was fixating on the center. The object was randomly selected from 12 (in the social-familiarity condition) or eight images (in the stable object-value condition, Fig. 1B). A fixed amount of reward was delivered after 2–4 sequential presentations. In this manner, neuronal responses would reflect long-term object-value memories rather than predicted rewards. We recorded the activity of putative medium spiny neurons during the passive-viewing task (22 from monkey W, 27 from monkey S, and 19 from monkey Z). Figure 2A shows the activity of a representative neuron in the STRt that responded to the facial images. The neuron showed a stronger response to the social familiarity (+) faces (top) than to the social familiarity (-) faces (middle; p < 0.05, Fisher’s test), although the responses to the social familiarity (+) faces varied. This neuron also showed a stronger response to the value (+) objects than to the value (-) objects (Fig. 2B, p = 0.01, Wilcoxon rank-sum test). Responses to valued objects also widely varied. These results are consistent with those of our previous studies, which showed that STRt neurons encode stable object values but are sensitive to particular objects (*5, 18*). The neuron with low visual selectivity also showed strong responses to the socially familiar faces and high-value objects (p < 0.05, p = 6.94 × 10^-4^; Fig. S2).

**Fig. 2.**
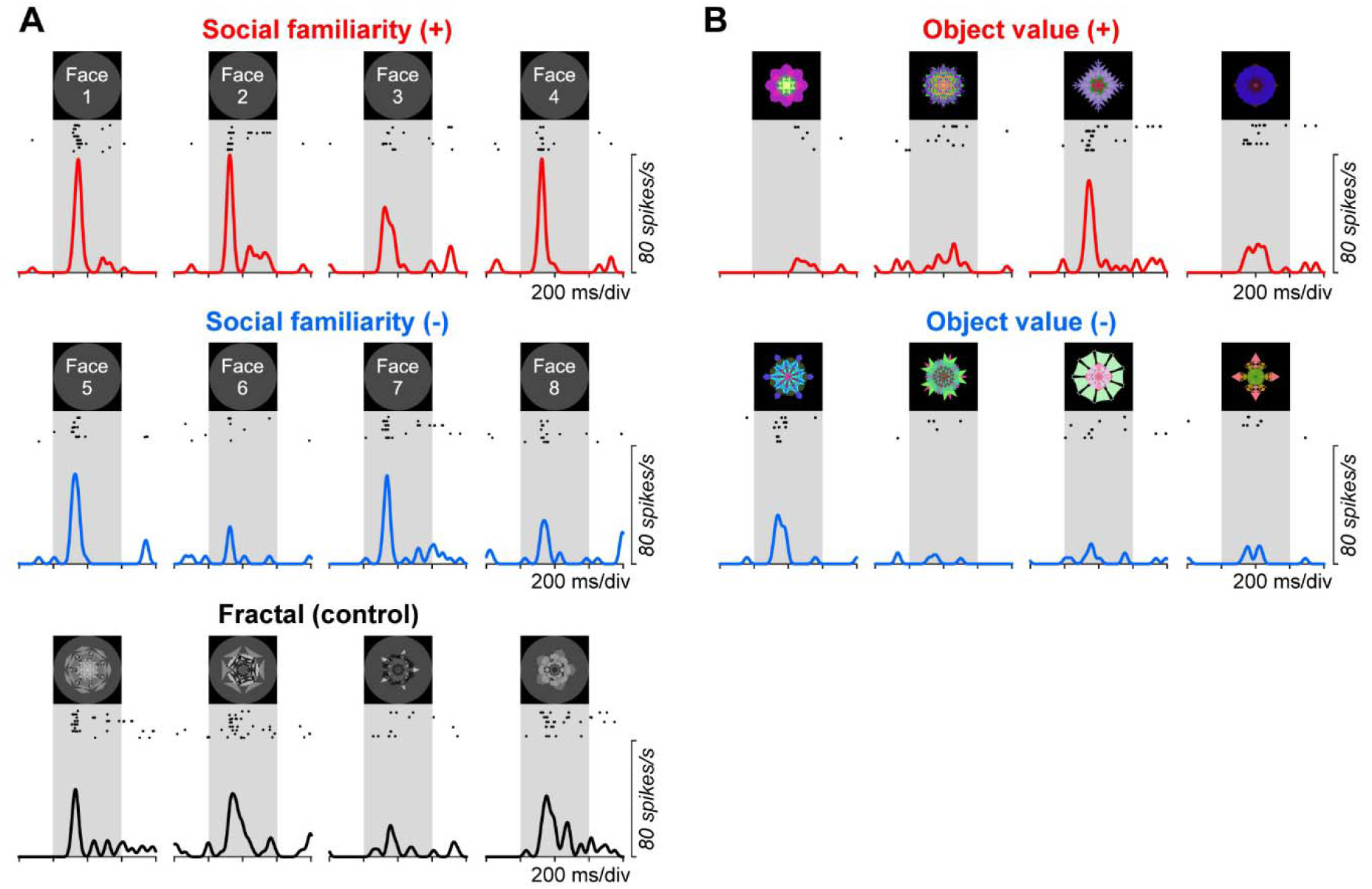
A representative face-responsive neuron in the striatum tail. **(A)** Neuronal response to the socially familiar (+) faces (top), socially familiar (-) faces (middle), and fractal objects (bottom) in the passive-viewing task. Face photographs were omitted. **(B)** Neuronal response to the value (+) objects (top) and value (-) objects (bottom). Same neuron as **(A)**.

First, we analyzed the responses during the social-familiarity condition to examine the property of facial responses of STRt neurons. Among 68 recorded neurons, 52 (75%) showed a significant visual response (Fig. 3A, p < 0.05 with Bonferroni correction, paired t-test). Fifty percent of neurons responded to the facial images (face-responsive neurons). We quantified the strength of social-familiarity coding for each neuron (modulation index, see Methods) and showed their distributions among all face-responsive neurons (Fig. 3B). The index was often away from 0.5, indicating that many neurons encoded social familiarity for facial images. A receiver operating characteristic (ROC) area of > 0.5 indicated that the neurons showed a stronger response to social familiarity (+) faces than to social familiarity (-) faces. In all, STRt neurons discriminated the socially familiar faces from the socially unfamiliar faces (p = 0.0051, unpaired t-test).

**Fig. 3.**
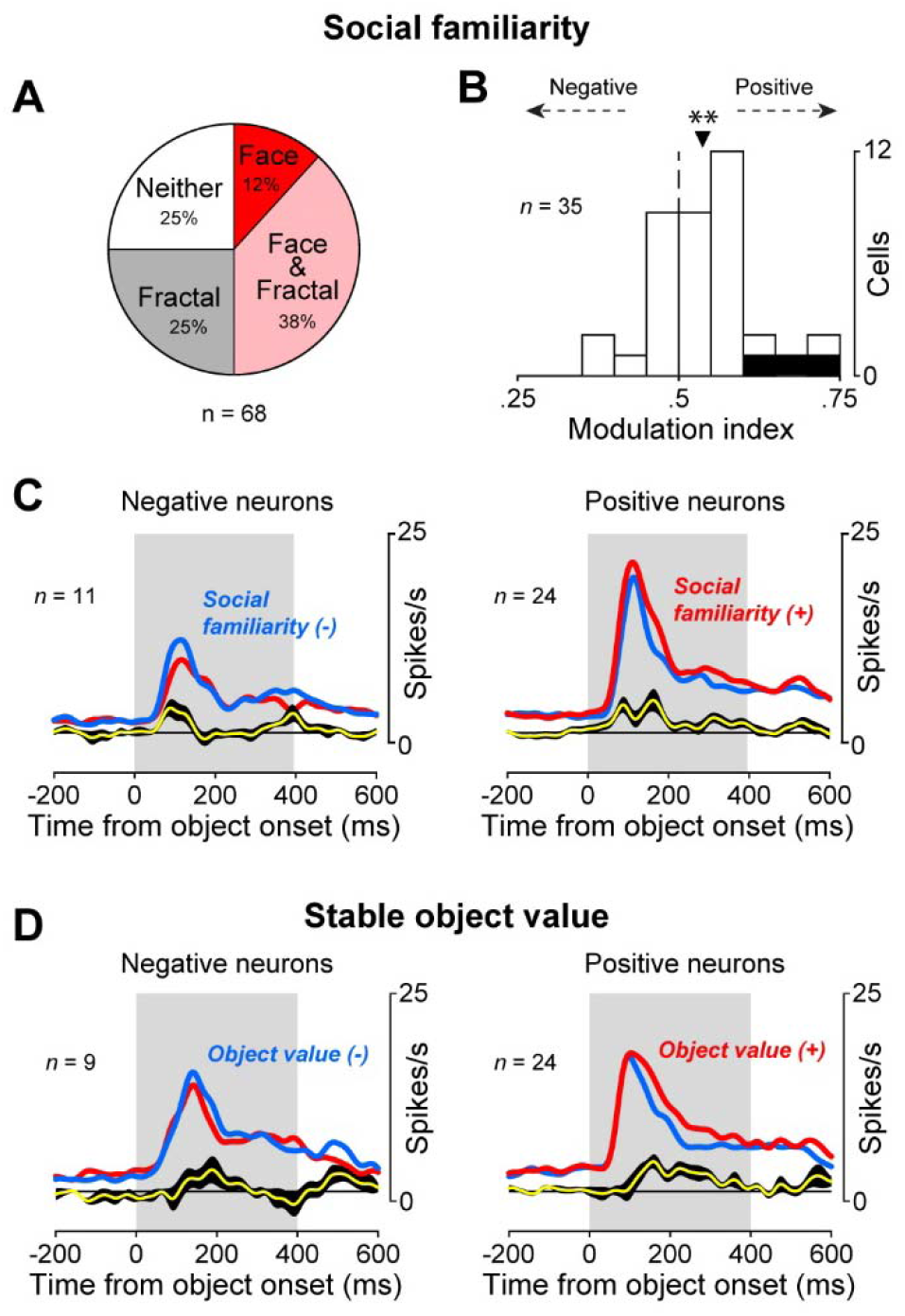
Face-responsive neurons encode social familiarity. **(A)** Proportions of visual neurons in the striatum tail categorized by the passive-viewing task. **(B)** Social familiarity modulation index of individual face responsive neurons in the striatum tail during the passive-viewing task. Filled bars indicate the neurons significantly modulated by social familiarity (p < 0.05, Fisher’s exact test). A value modulation index score of 1.0 signifies that the neuronal response to familiar faces is always stronger than that to unfamiliar faces; 0.0 signifies the opposite response pattern. The triangle indicates the population average of the modulation index. Asterisks indicate a significant difference from 0.5 (**P < 0.01, paired t-test). **(C)** Population neuronal activity in the social-familiarity condition; negative social-familiarity coding neurons (negative neurons, left) and positive social-familiarity coding neurons (positive neurons, right). Red and blue lines indicate the averaged neuronal responses to socially familiar faces and unfamiliar faces, respectively. The yellow line indicates the difference in response to the socially familiar faces and unfamiliar faces (mean ± SEM). **(D)** Population neuronal activity in the stable object-value condition; negative-value coding neurons (negative neurons, left) and positive-value coding neurons (positive neurons, right). Red and blue lines indicate the averaged neuronal responses to value (+) and (-) objects, respectively. The yellow line indicates the difference in response to value (+) and (-) objects (mean ± standard error of the mean).

Our previous study revealed that there are two groups of neurons in the STRt: a) positive-value coding and b) negative-value coding neurons (*4, 5, 7*). Accordingly, we analyzed the time course of population activity of these face-responsive neurons according to each face-coding type (positive or negative) (Fig. 3C). Social-familiarity coding started approximately 50 ms after the appearance of the image in both groups. These results suggested that STRt neurons quickly detect socially familiar persons. To examine whether these neurons also encode stable object values, we analyzed the activity of 33 face-responsive neurons in the STRt during the passive-viewing task with long-term reward-associated objects. Figure 3D shows the population activity of negative and positive-value coding neurons (the neuron types were defined by the response to socially familiar and unfamiliar faces). The value modulation index of individual STRt neurons was biased toward 1.0 (p = 0.023, unpaired t-test), which would indicate the absolute preference for positive value objects. Interestingly, the detection of social familiarity preceded that of stable object value in both coding types.

From the above, we demonstrated the encoding of two types of information in STRt neurons: a) social familiarity, b) object value. We then asked whether these signals are encoded by different or the same neurons in the STRt. Figure 4 shows that the strengths of social familiarity bias and object value bias were significantly correlated (Pearson’s r = 0.41, p = 0.02). These results indicate that social information and value information are co-encoded in the STRt and that these signals are mediated by the same neuronal mechanism.

**Fig. 4.**
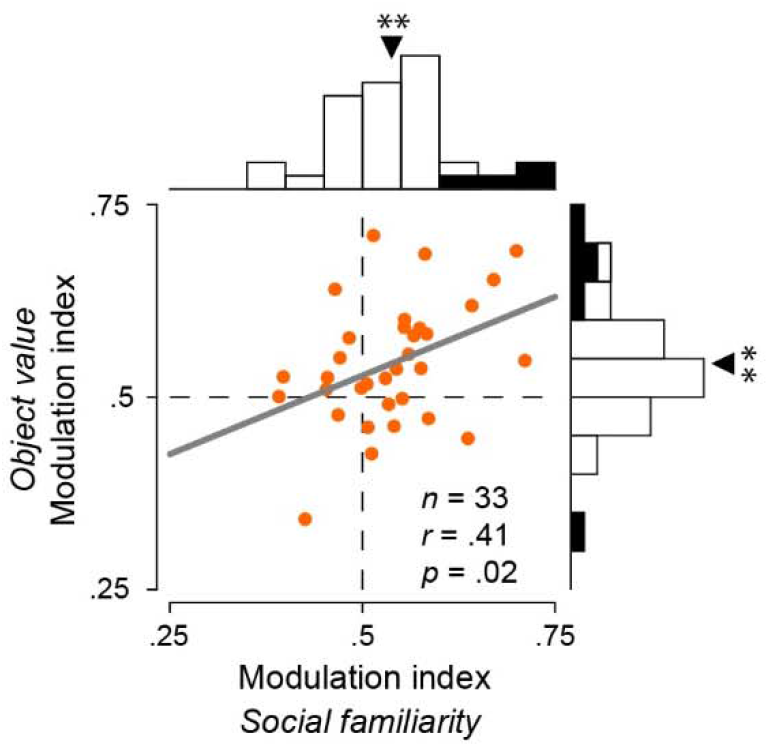
Correlation between social familiarity and stable object value. The correlation between modulation indices of social familiarity and stable object value. Numbers indicate Pearson’s correlation coefficient and P value. Note that the strength of object-value coding is significantly correlated with that of social-familiarity coding.

## Discussion

Although the basal ganglia play a key role in reward value-guided behavior based on experience (*19*), it is less clear whether they are involved in social behavior. This was the first study to report neuronal signals in the basal ganglia in relation to face recognition. We first found that many STRt neurons responded to facial images (Fig. 2). The response to socially familiar faces was stronger than that to socially unfamiliar faces (Fig. 3). Then, we compared the response to objects based on long-term memory with the response to faces based on social familiarity. The modulations of social familiarity and stable object value were positively correlated (Fig. 4). Our results suggest that the posterior basal ganglia circuit co-encodes social familiarity and stable object value.

Research on human face perception is a major neuroscience topic. Multiple face-responsive areas have been found in the primate temporal cortex (*2, 3*). Microstimulation of face-selective sites in the inferior temporal cortex (IT) around the image presentation period biased monkey decision toward faces during a face categorization task (*20*). These findings suggest that the temporal cortex plays a key role in face identification. Furthermore, the TE and temporal pole show strong responses to personally familiar faces (*21-23*). These studies raise the question on which neuronal circuit encodes socially familiar faces? In this study, we observed that neurons in the STRt, which is robustly connected to the TE, showed face selectivity (Fig. 2, (*11, 12, 24, 25*)). The basal ganglia encode object values based on gradual change in synaptic weight, which is modulated by the reward signal of dopamine (*26-28*). Especially, our previous studies suggested that short-term and long-term value memories are mediated by the anterior and posterior parts of the basal ganglia circuit, respectively (*29*). It is possible that the basal ganglia circuit for social familiarity is similar to the circuit for stable object value because the subjects recognized persons based on emotional and reward experiences, social familiarity is based on long-term experience and long-term value memory, and the same STRt neurons are involved in both processes. Indeed, we found a positive correlation between social familiarity and stable object value modulations (Fig. 4).

Humans can detect familiar faces more quickly and accurately than unfamiliar ones (*30, 31*). Our recent studies showed that the visual responses in STRt neurons lead to quick detection of high-value objects (7). STRt responses to high-valued objects are accomplished with high object selectivity (*5, 18*). Similar high object selectivity was also found in the STRt response to facial images (Fig. 2). This may be based on numerous inputs from many neurons in the temporal cortex to single STRt neurons. Moreover, in the STRt, neurons are divided into two groups: positive neurons strongly responsive to value (+) objects and social familiarity (+) faces and negative neurons strongly responsive to value (-) objects and social familiarity (-) faces (Fig. 3). Previous studies have suggested that these preferences are related to the neuronal process of oculomotor control. The STRt projects to two local regions in the basal ganglia [caudal–dorsal–lateral part of the substantia nigra pars reticulate (cdlSNr) and caudal–ventral part of the globus pallidus externus (cvGPe); (*8, 10*)]. The inhibitory connection to the cdlSNr, which acts as the direct pathway, conveys the high-valued object-dominant signal and inhibits cdlSNr neurons, leading to the disinhibition of neurons in the superior colliculus (SC) to facilitate saccades toward high-valued objects (*32, 33*). The inhibitory connection to the cvGPe, which acts as the indirect pathway, conveys the low-valued object-dominant signal and inhibits cvGPe neurons (*8*). As cvGPe neurons have inhibitory connections to cdlSNr neurons (*34-36*), the output of the CDt disinhibits cdlSNr neurons, leading to enhanced inhibition of SC neurons to suppress saccades toward low-valued objects (*8, 33*). The same mechanism may be underlying the facilitation and suppression of saccades toward facial images.

In this study, we used human facial images to test the neuronal representation of social familiarity in monkeys because, in the laboratory, human–monkey interaction is relatively simpler and clearer than monkey–monkey interaction. Close interaction between species is commonly encountered between owners and companion animals. Our results comprise evidence regarding the neuronal mechanism of these interactions. It is unclear whether the monkeys behaviorally distinguished the socially familiar facial images from the unfamiliar ones used in the present study (Fig. 1). In real-world settings, person identification relies both on facial and on physical-characteristic, voice, and odor cues. Therefore, grayscale facial images may not contain sufficient information to identify individuals. However, a previous study reported that it is possible to identify a familiar face from a poor-quality image (*37*). Monkeys may be able to distinguish facial images of familiar versus unfamiliar persons without clearly identifying these persons.

Previous studies have reported that patients in the early stage of Huntington’s disease, who have degeneration of CDt neurons (*15*), show deficits in face recognition and discrimination (*16*). These reports are consistent with our findings in that the STRt is involved in face representation. In contrast, patients with Parkinson’s disease, which is caused by dopamine deficits, have normal face-detection abilities despite having impaired facial-emotion recognition ability (*38*). The loss of dopamine typically leads to dysfunction of the anterior part of the striatum, including the nucleus accumbens (*39, 40*). Notably, people with autism, who have impaired social communication abilities, show no deficits in the recognition of familiar faces (*41*). This report supports our conclusions that the same mechanism is responsible for the encoding of socially familiar faces and stable object values in the STRt, independent of the circuit of social behavior.

## Materials and Methods

### Animal Model

Three adult male rhesus monkeys (*<u>Macaca mulatta</u>*, 8–11 kg, 6–8 years old) were used for all experiments. All procedures for animal care and experimentation were approved by the Animal Care and Use Committee of the National Eye Institute (Bethesda, USA) and complied with the Public Health Service Policy on the humane care and use of laboratory animals.

### General Procedures

We implanted a plastic head holder, eye coil, and plastic recording chamber under general anesthesia and sterile surgical conditions. After the monkeys fully recovered from surgery, we started training them in the oculomotor tasks. Several procedures, including the surgery, behavioral task, and statistical analysis, were identical between this and our previous studies (*5, 42*) .

### Behavioral Procedure

The behavioral procedure was controlled by a C++ based real-time experimentation data acquisition system (Blip: available at http://www.robilis.com/blip/). The monkey sat in a primate chair, facing a frontoparallel screen in a sound-attenuated and electrically shielded room. Visual stimuli generated by an active matrix liquid crystal display projector (PJ550, ViewSonic) were rear projected on the screen. Facial images (10° × 10°) were provided by a member of the laboratory. Four persons [experimenters and animal caretakers, social familiarity (+)] were involved in monkey daily care and were familiar with each monkey. The other four persons [social familiarity (-)] had not met the subjects. We created the fractal objects (□10° × 10°) using fractal geometry (*18*).

### Passive-Viewing Task (Social-Familiarity Coding Test)

The purpose of this task was to examine the response property of the striatal neurons to facial images. We used grayscale images of faces (socially familiar and unfamiliar persons’ faces) and of fractal objects (no value association) to test the neuronal response to facial images (Figure 1A). In this task, two to four facial images (facial images or fractal objects) were sequentially presented in the receptive field (presentation time, 400 ms; interval, 400 ms) while the monkey fixated on a central white dot (Figure 1D). A fixed liquid reward (0.2 ml) was delivered 300 ms after the last presentation. The reward was thus not contingently associated with any facial image. Each image was presented at least seven times in each session.

### Stable Object Value Task

We used the stable object value procedure to examine the long-term effect of object value learning (*5, 6*). Eight fractal objects were divided into four high-valued objects [object value (+)] associated with a large reward (0.3 ml) and four low-valued ones [object value (-)] associated with a small reward (0.1 ml) (Figure 1B). The monkey learned each object value by making a saccade to each object followed by large or small reward delivery (Figure S1C). One learning session consisted of 80 trials. The monkey was trained for each set of objects with one learning session each day. The same sets of objects were repeatedly used across at least five learning sessions.

### Free-Viewing Task

The free-viewing task was used as a behavioral test to examine monkey preference for value (+) or value (-) objects (*5, 6*). After the monkey fixated on a central white dot for 300 ms, four objects were simultaneously presented in four symmetric positions (15° from center) (Figure 1C). The four objects were pseudo-randomly selected from eight objects used in the stable object value task. The monkey was free to look at the objects for 2 s without a reward outcome. After a blank period (500 ms), another white dot was presented at one of eight positions. When the monkey made a saccade to it, a fixed reward was delivered (0.2 ml). Each object was presented at least 10 times in each session.

### Passive-Viewing Task (Stable Object Value-Coding Test)

Once the monkey showed significant behavioral preference for value (+) objects in the free-viewing task, we tested neuronal responses to the objects with the passive-viewing task (*5, 6*). This was the same as the passive-viewing task for the social-familiarity coding test except for the presented images. Two to four objects used in the stable object value task were sequentially presented in the receptive field while the monkey fixated on a central white dot. A fixed liquid reward (0.2 ml) was delivered after the last presentation. Therefore, each object was no longer associated with a large or small reward in this task. We usually used two to three sets of well learned objects.

The behavioral test (free-viewing task) and the neuronal test (passive-viewing task) were conducted on different days. To remind the monkey of the object value memory, we conducted the stable value task at least once per 60 days after the initial learning.

### Recording Procedure

Based on a stereotaxic atlas, a recording chamber was placed over the parietal cortex, tilted laterally by 25° (monkey W and Z) or 0° (monkey S) and aimed at the STRt. Magnetic resonance images (4.7 T; Bruker) were then obtained along the direction of the recording chamber, which was visualized by filling a recording grid with gadolinium.

For single-neuron recording, a tungsten electrode (Alpha Omega Engineering or FHC) was lowered into the striatum through a guide tube using a micromanipulator (MO-97S; Narishige). The recording site was determined using a grid system, which allowed electrode penetrations at every 1 mm. We amplified and filtered (0.3–10 kHz; Model 1800, A-M Systems; Model MDA-4I, BAK) the signals obtained from the electrodes and collected at 1 kHz. Single neurons were isolated online using custom voltage–time window discriminator software (Blip).

### Statistical Analysis

We defined the putamen tail as the region 0–3.5 mm from the ventral edge of the putamen and the cdPUT as the region above it (Kunimatsu et al., 2019; Kunimatsu et al., 2021). Because the putamen tail and CDt share the same anatomical pathway (*10, 12*) and showed similar long-term value coding in a previous study (*5*), we combined neurons in these areas as STRt neurons. The offline analyses were performed using Matlab (MathWorks). We defined visual neurons as neurons showing a significant difference in activity between the baseline period (400 ms before object onset) and visual period (50– 350 ms after object onset; paired t test, p < 0.05 with Bonferroni correction). We considered “face-responsive” and “object-responsive” neurons showing a significant response to the faces and fractal objects, respectively. The time course of neuronal activity for each condition is shown after smoothing with a Gaussian kernel (σ = 15 ms). We only used the data in correct trials for the behavioral and neuronal analyses.

To examine neuronal discrimination, we measured the magnitude of the response to each visual stimulus by counting the numbers of spikes fired (on single trials) within a test window (50–350 ms after visual stimulus onset). A value modulation index was defined as the area under the ROC curve based on the response magnitude of the neuron to facial images [social familiarity (+) vs. (-)] or fractal objects [object value (+) vs. (-)].

## Acknowledgments

We are grateful to R.H. Wurtz, C. Quaia, L. Wang, K. McAlonan, D. Parker, O. Mohammed, and D.M. Prebilic for providing facial images. We thank D. McMahon for manuscript-writing assistance and D. Parker, V. McLean, D.M. Prebilic, O. Mohammed, G. Tansey, A.M. Nichols, T.W. Ruffner, and A.V. Hays for technical assistance.

## Author contributions

Study design: J.K. and O.H.

Experiments: J.K and H.A.

Data analysis: J.K.

Writing: all authors

## Data and materials availability

The data that support the findings of this study are available from the corresponding author upon reasonable request.

