## Supplemental figures for "Neuronal mechanism of the encoding of socially familiar faces in the striatum tail"

**This PDF file includes:**

Figs. S1 to S2

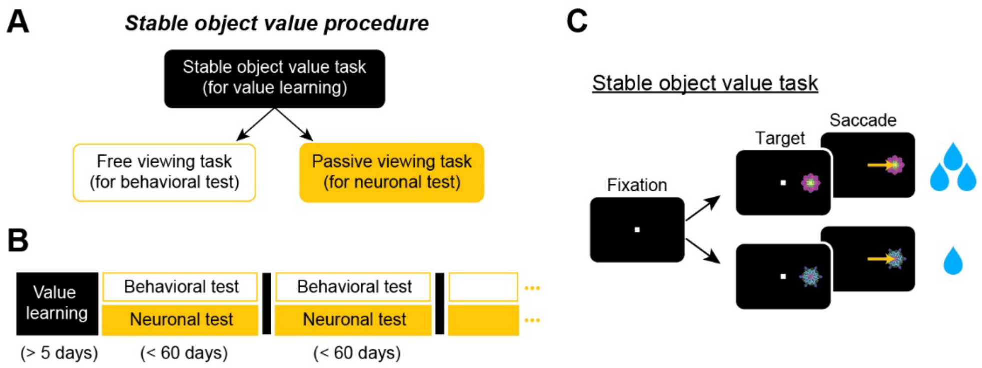


**Fig. S1. Stable object value procedure.**

(**A**) The stable object value procedure consisted of two steps: learning and test. After across-day learning (stable object value task; **C**), its learning effect on the subject's behavior was tested with the free-viewing task (Fig. 1C) and the neuronal response with the passive viewing task (Fig. 1D).

(**B**) Schedule of learning and test phases: behavioral and neuronal tests were conducted after the learning sessions (> 5 days). The learning session was conducted at least once per 60 days.

(**C**) Stable object value procedure. After the monkey made a saccade to the presented object, a large or small reward was delivered. Four objects were associated with a large reward [object value (+)] and four objects were associated with a small reward [object value (-)]. The reward amount was fixed for each object.

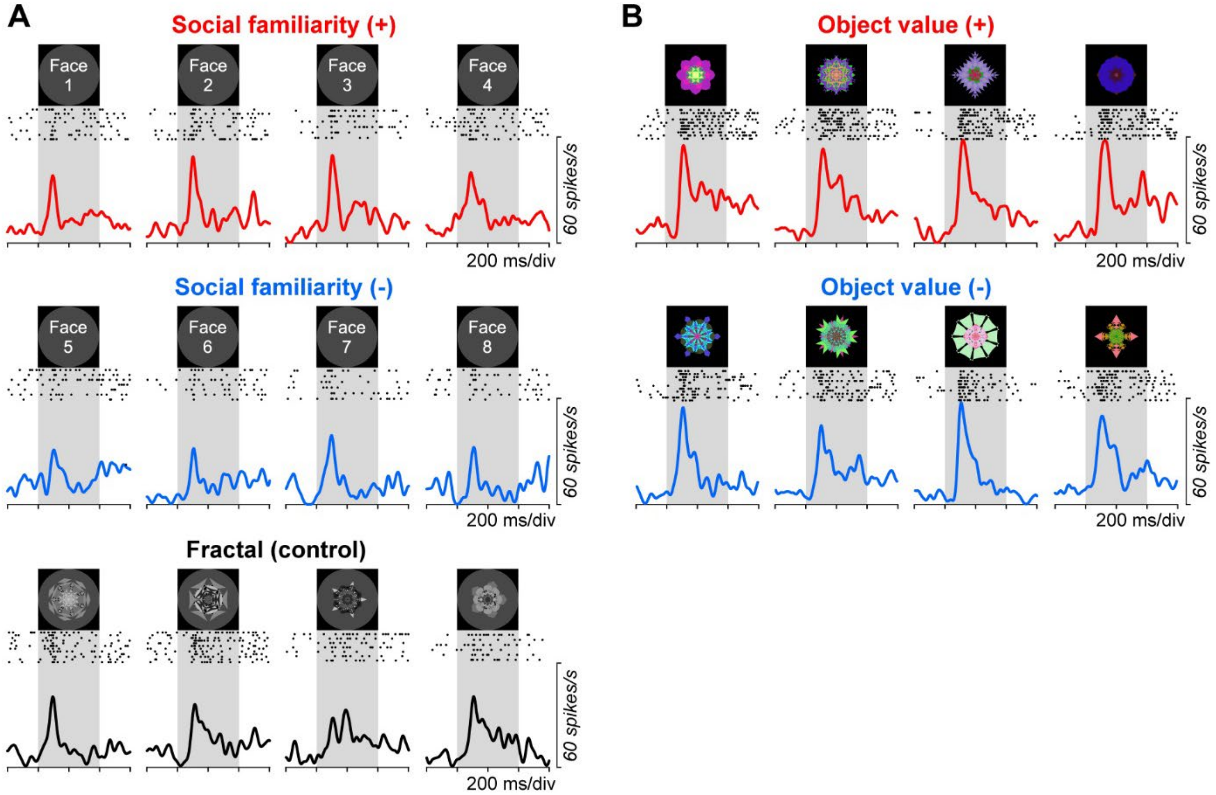


**Fig. S2. Another representative face-responsive neuron in the striatum tail.**
This neuron shows clear social familiarity and stable object value but less selectivity to each face and each object. Face photographs were omitted. The format is the same as that in Fig. 2.
